# Bio-Inspired Molecular Bridging in a Hybrid Perovskite Leads to Enhanced Stability and Tunable Properties

**DOI:** 10.1101/2020.08.03.233783

**Authors:** Arad Lang, Iryna Polishchuk, Eva Seknazi, Jochen Feldmann, Alexander Katsman, Boaz Pokroy

## Abstract

Hybrid organic-inorganic halide perovskites demonstrate high potential in several applications such as solar cells, field-effect transistors, light-emitting diodes and more. However, the main drawback which limits their use in such applications is their low stability in humid conditions. In this paper we implement one of Nature’s strategies found in bio-crystals in order to improve the stability of the hybrid perovskite methylammonium lead bromide (MAPbBr_3_) in water, as well as to tune its structure, optical and thermal properties. This was achieved, for the first time, by the incorporation of amino acids into the lattice of MAPbBr_3_. The amino acid lysine, which possesses two NH^3+^ groups, is incorporated into the hybrid unit cell, by substituting two methylammonium ions and serves as a “molecular bridge”. This incorporation induces a decrease in the lattice parameter of the host, accompanied with an increase in the band gap and noticeable changes in its morphology. Furthermore, we observed an increase in thermal expansion coefficient and a shift of the phase transformation temperature of the hybrid crystal. The level of amino acid incorporation depends on the conditions of crystallization, which also influence the extent of MAPbBr_3_ band gap changes. Notably, lysine incorporation strongly increases the perovskite stability in water. This study demonstrates the unique and promising approach to tune the properties and stability of hybrid perovskites via this novel bio-inspired route.

## Introduction

In recent years, hybrid organic-inorganic perovskites (HOIPs)^[1–3]^ have attracted widespread attention due to their utility in various applications, such as photovoltaics,^[4–8]^ light emitting diodes (LEDs),^[9–11]^ and photodetectors.^[12–14]^ These HOIPs have been widely studied considering their exceptional conversion efficiency,^[8,15,16]^ together with easy bulk synthesis,^[17–19]^ device fabrication,^[4,20]^ and band gap engineering techniques.^[21]^

One major setback which hinders HOIPs from being commercially used is their low stability.^[22,23]^ This family of materials is chemically active and thermally unstable.^[24,25]^ They tend to degrade and decompose while reacting with water (humidity),^[26,27]^ oxygen^[28]^ and even light itself.^[29]^ In order to overcome this difficulty, many attempts have been done, among them adding organic^[30–32]^ or oxide^[33]^ protective layers, ion substitutions^[34,35]^ and morphology control.^[36–39]^ Given the remarkable potential of these systems and the importance of novel pathways to tune the physical properties of HOIPs, as well as improve their stability, we were interested in the possibility of a Nature-inspired tuning strategy: via the incorporation of amino acids into HOIPs’ crystal lattice and its effects on the crystal structure and properties.

Biominerals, owing to their superior mechanical properties, have been a source of inspiration to scientists for many years.^[40–48]^ Common examples are mollusk shells, which are remarkably hard and tough,^[49,50]^ although they are composed of calcium carbonate (CaCO_3_), a brittle ceramic. Such enhancement of mechanical properties originates from several factors, which include both intra- and inter-crystalline organics along with unique hierarchical structures.^[49,51–53]^

Interaction between the inorganic matrix and the organic inclusions (usually peptides or polysaccharides) in biominerals is complex and is not yet fully understood.^[54,55]^ Calcite, the most thermodynamically stable polymorph of CaCO_3_,^[56]^ was nevertheless shown to be capable of incorporating proteins,^[55,57]^ polymer nanoparticles,^[58,59]^ dyes,^[60]^ inorganic nanoparticles,^[61–63]^ graphene oxide,^[64]^ anticancer drugs,^[65]^ gels^[66,67]^ and individual amino acids.^[68,69]^ Incorporation of amino acids induces expansion of the unit cell (*i*.*e*., an increase in the lattice parameters, both *a* and *c*), and is accompanied by significant changes in the crystal morphology.^[68,69]^ Furthermore, the incorporated amino acids enhance the hardness of the synthetic calcite single crystals, as they create internal stresses and act as precipitates in a second phase-hardening mechanism.^[69]^

The ability to incorporate amino acids does not belong exclusively to calcite. Zinc oxide (ZnO, wurtzite) and copper oxide (Cu_2_O, cuprite) were also shown to be capable of hosting individual amino acids.^[70,71]^ For ZnO the effect of the incorporation is similar to that of calcite; in both cases the unit cell expands, and there are changes in crystal morphology.^[70]^ In contrast, incorporation of amino acids into Cu_2_O causes shrinkage of its unit cell, owing to substitution of the Cu^+2^ by the amine moiety of the amino acid.^[71]^ Interestingly, this incorporation induces an increase in the optical band gaps of both of the inorganic semiconductors.^[70–72]^ This increase results from a quantum-size-like effect, in which localization of charge carriers is induced by dispersion of the insulating amino acids within the semiconducting matrix.^[72]^

In this study, our material of choice from the HOIP family is methylammonium lead bromide (MAPbBr_3_).^[10,11,73–76]^ It possesses the classic cubic perovskite ABX_3_ structure, with CH_3_NH_3+_ (MA^+^) cations in its A-sites, which are asymmetric and randomly aligned.^[77,78]^ Incorporation of amino acids into the lattice structure of HOIPs has never before been performed. We therefore aimed to investigate whether such incorporation is feasible in this system, and if so, to determine its effects on the crystal structure and properties.

## Results and Discussion

We synthesized MAPbBr_3_ crystals from solution by utilizing the method described by Poglitch & Weber.^[17]^ In a typical synthesis, a Pb_+2_ solution was prepared by dissolving lead acetate trihydrate (Pb(OAc)_2_) in concentrated hydrobromic acid (HBr). The same amount of all the 20 common L-amino acids were added to the Pb^+2^ solution. Precipitation of MAPbBr_3_ was activated by the addition of an equimolar volume of MA^+^ solution to the Pb^+2^ solution (for more details see Experimental Section). Crystallization was achieved via two different routes: fast synthesis, in which the MA^+^ solution was added to the Pb^+2^ solution at room temperature while stirring, and slow synthesis, in which the MA^+^ solution was added to a hot Pb^+2^ solution (in an oil bath at 95°C) and the mixture was cooled down in air. In both cases, the resulting orange-hued precipitates were filtered, washed with acetone, and dried in air.

As a first analysis of the precipitated crystals, we characterized their structure utilizing high-resolution powder X-ray diffraction (HR-PXRD). By applying Rietveld refinement^[79]^ on the full diffraction patterns we were able to deduce the lattice parameters of the crystals with high precision. When we compared the crystal structure of crystals formed with and without amino acids in the crystallization solution, lattice contraction could be seen in the former case (**Figure 1)**. Exceptions in this regard were Asn, Gln, and Trp. Asn and Gln were excluded because of spontaneous decomposition of the amide group in the acidic solution, and Trp was not soluble. From **Figure 1** (and **Table S1**) it is evident that by far the highest distortions are induced by Lys, indicating that this amino acid induces the most pronounced effect.^[68,70,71]^ Moreover, it is clear that incorporation of the other amino acids was lower, and was at approximately the same level in both growing routes. Since the effect of Lys on the MAPbBr_3_ lattice was significantly higher, especially via the slow growth procedure, we focused in this work on the study of Lys incorporation. MAPbBr_3_ samples were grown from solutions containing different Lys concentrations: 0.00, 0.02, 0.04, 0.06 and 0.08 g ml^-1^.

**Figure 1.**
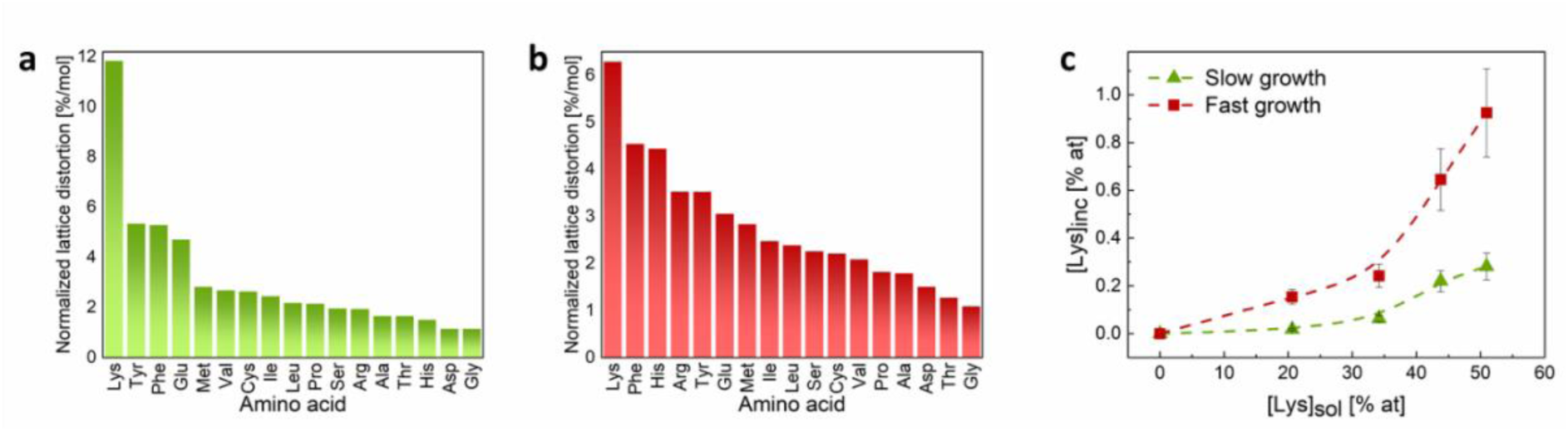
Incorporation of amino acids into MAPbBr3 lattice. Normalized lattice distortions in MAPbBr_3_ crystals induced during their growth in the presence of different amino acids: (a) slow growth; (b) fast growth. The normalized distortions are the measured distortions (using HR-PXRD) divided by the amount of amino acid molecules in solution. (c) Concentration of incorporated Lys (measured by amino acid analysis) vs. its concentration in solution under conditions of slow and fast growth.

After dissolving the crystals, we used amino acid analysis (AAA) to estimate the correlation between the concentration of incorporated Lys ([Lys]_inc_) and its initial concentration in solution ([Lys]_sol_) **(Figure 1c)**. As expected, increasing the concentration of Lys in the Pb^+2^ solution (prior to addition of MA^+^ to the solution) resulted in a larger amount of incorporated Lys. It is also apparent from **Figure 1c** that the fast crystallization led to higher incorporation levels than those of the slow process, reaching a maximum of almost 1 at.%. In fact, the level of Lys incorporation into the lattice of MAPbBr_3_ was of the same order of magnitude as that of Asp incorporated into calcite^[69]^ and of Asn incorporated into Cu_2_O.^[71]^ More interestingly, HR-PXRD analysis revealed that the incorporation of Lys into MAPbBr_3_ crystals induces a shift of the diffraction peaks to higher 2θ values (**Figure 2a**), which—according to Bragg’s law^[80]^— correspond to smaller inter-planar spacings. Similar lattice shrinkage rather than expansion was observed upon incorporation of amino acids into Cu_2_O semiconductor.^[71]^ Plotting the lattice contraction as a function of incorporation levels **(Figure 2b, Table S2)** clearly demonstrates that for both growth regimes, fast and slow, the higher the level of Lys incorporation, the higher the induced lattice shrinkage.

**Figure 2.**
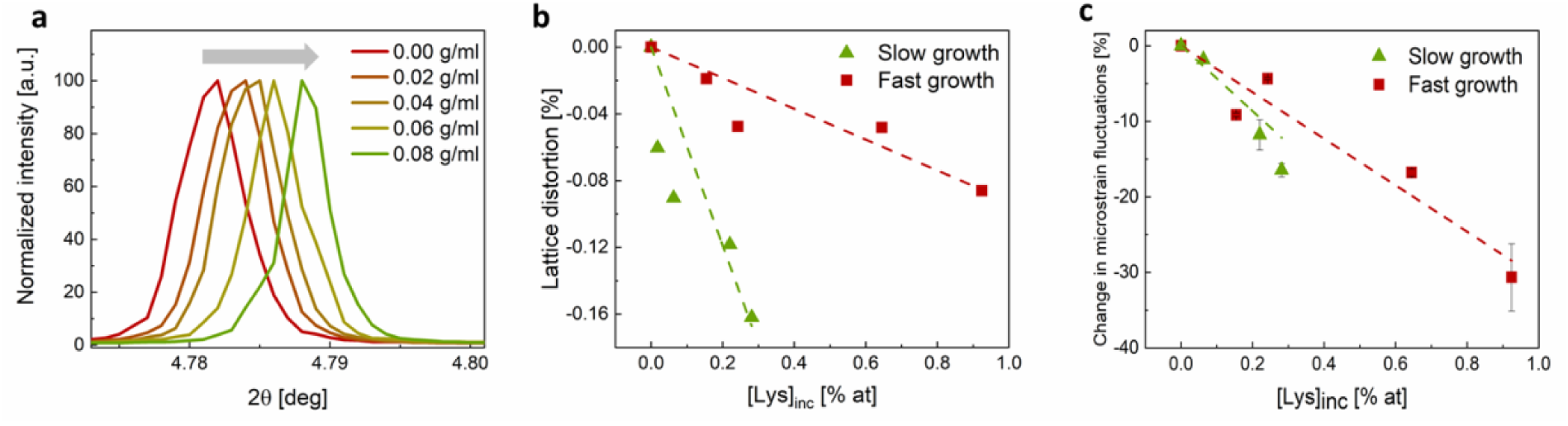
Lattice distortion induced by Lys incorporation. (a) The {100} diffraction peak of MAPbBr_3_ grown in the presence of different amounts of Lys (λ=0.49599 Å). (b) Lattice distortions and (c) relative change in microstrain fluctuations of MAPbBr_3_ vs. concentration of the incorporated Lys.

We also fitted a pseuso-Voight function to the {100} diffraction peak of each sample and calculated the average grain size and microstrain fluctuations based on the full width at half maximum (FWHM, see Table S3).^[81]^ As shown in **Figure 2**c, the microstrain fluctuations decreased with increasing amounts of Lys in the crystals. This trend differs from the widespread phenomenon in which the incorporation of organics is known to cause an increase in FWHM^[68,70]^ in biogenic crystals as well as in amino acid incorporated synthetic calcite.

The incorporation of about 1 at.% of an amino acid into MAPbBr_3_ has the potential to alter the thermal properties of the host crystal. To verify this assumption, the lattice parameter of each sample was measured via HR-PXRD coupled with in-situ cooling at different temperatures (290°K, 275°K, 250°K, and 230°K – see **Figure S4**). Cooling was achieved using a cold nitrogen gas blower. The lattice expansion coefficient was calculated according to equation (2):

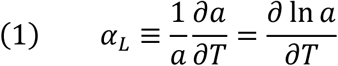

where *a* is the measured lattice parameter. By plotting the slope of ln(*a*) vs. the absolute temperature, the thermal expansion coefficient can be extracted.

In the case of Lys incorporation, a noticable effect on the thermal properties of MAPbBr_3_ was observed, namely a decrease in the thermal expansion coefficient **(Figure 3**a) with increasing Lys concentration. For changes in lattice parameter, the relation between the change in thermal expansion coefficient and lattice distortion can be estimated as:

**Figure 3.**
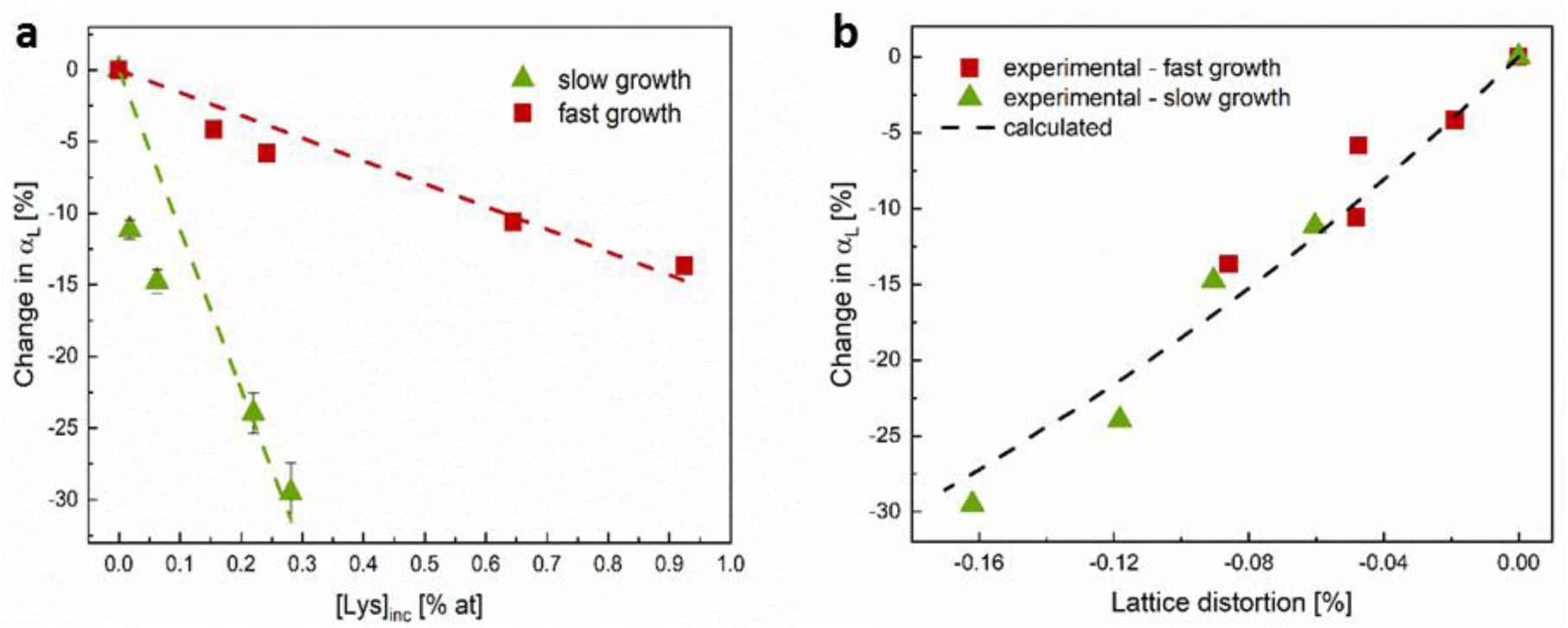
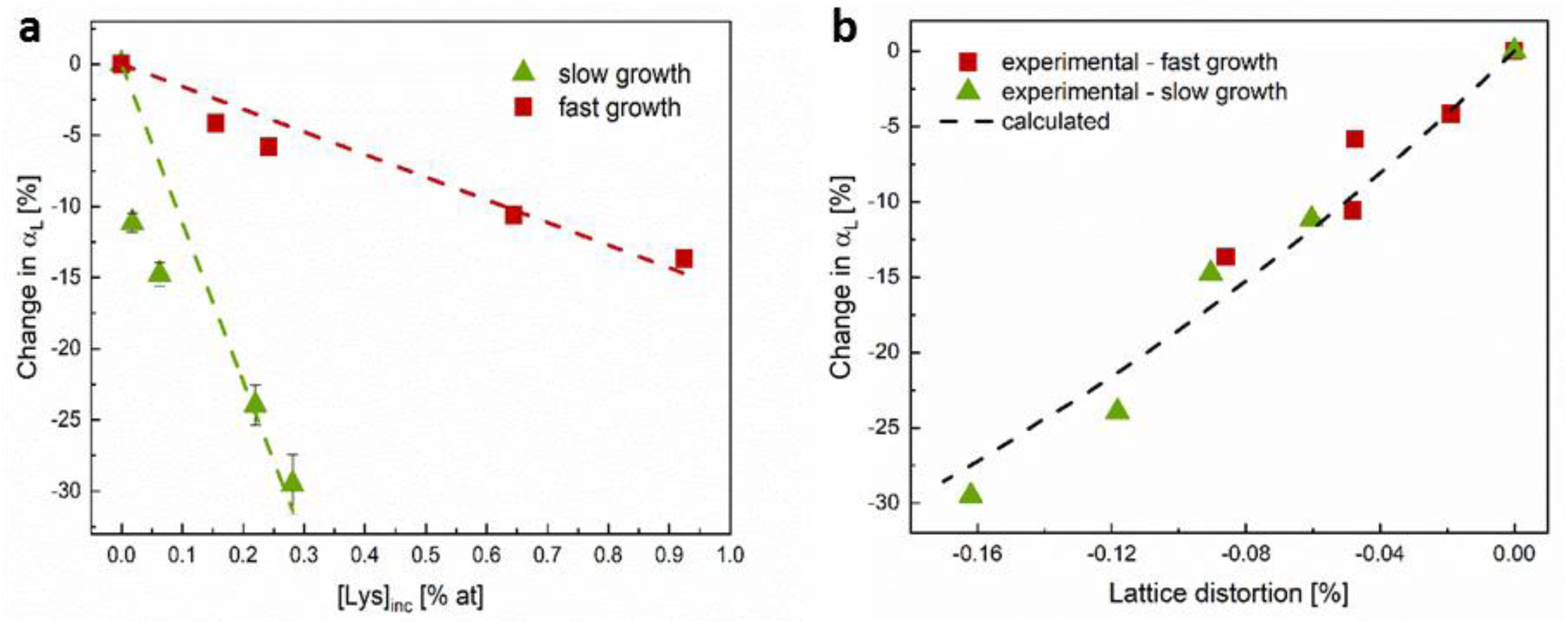
Change in thermal expansion coefficient. Change in MAPbBr_3_ thermal expansion coefficient *α*_*L*_ vs. (a) concentration of incorporated Lys, and (b) lattice distortion. The dashed line in (b) corresponds to equation (3).

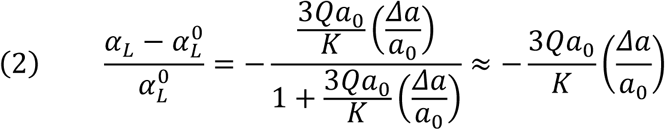

Where *Q* and *K* are coefficients of the power series expansion of the interatomic potential, and *Δa*/*a*_0_ is the (negative) lattice distortion (for more details, see supporting information). Accordong to equation (3), the change in thermal expansion coefficient is proportional to the lattice distortion. This result can explain both the trend and the differene between slow- and fast-grown MAPbBr_3_, as presented in

MAPbBr_3_ is known to undergo a cubic-to-tetragonal (C-T) phase transformation (from 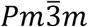 to *I*4/*mcm* space group) during cooling, at a temperature of about −36°C.^[17]^ In light of this, and since Lys incorporation was found here to affect the thermal expansion properties of MAPbBr_3,_ we also studied the effect of incorporated Lys on the C-T phase trasformation. To this end, using DSC, we evaluated the phase transformation temperature of a control sample as well as of Lys-incorporated samples. Each sample was scanned from room temperature to −80°C and back, at a rate of 5°C/min. The transformation temperature was taken as the temperature at the onset of the phase transformation peak (see **Figure S5a**).

Interestingly, we found that the temperature of the C-T phase transformation was indeed altered upon the incorporation of Lys **(Figure 4**a). Relative to the control sample the transformation temperature of the Lys-incorporated samples decreased during heating and increased during the cooling cycle. The temperature of transformation of pure MAPbBr_3_ is in good agreement with a previous report.^[17]^ It is clear, moreover, that this trend intensifies as the Lys incorporation level increases. Given that the incorporation occurs while MAPbBr_3_ is crystallizing in its cubic form (above the transformation temperature), this finding was the opposite of what we would expect from a standard colligative property^[82]^. One can suggest that the Lys incorporation inhibits the C-T transition in both directions by affecting the rotation of PbBr^6^ octahedra.^[83]^ Lys incorporation hinders this rotation, hence inducing a spreading of the phase transformation peak as was measured with DSC (**Figure 4b**) (for more details, see supporting information).

**Figure 4.**
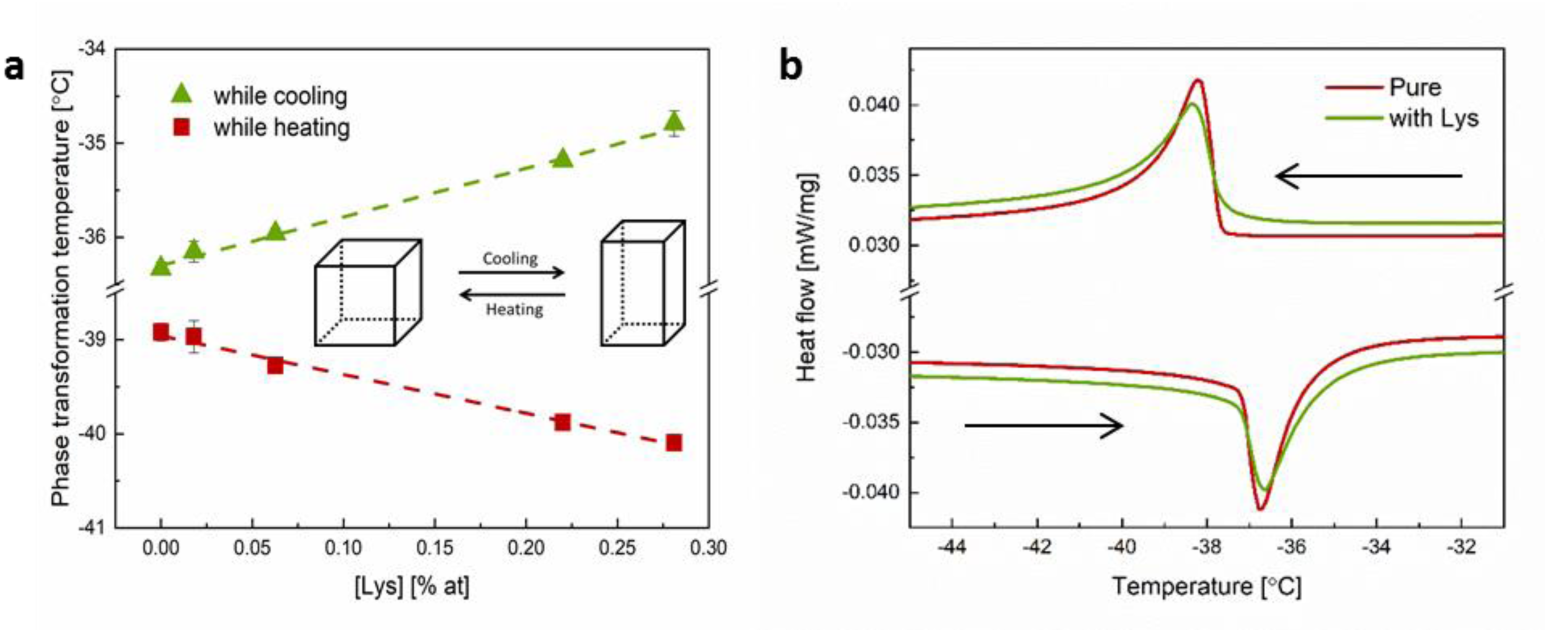
Change in transformation temperature. (a) Phase transformation temperature of slow-grown MAPbBr_3_ vs. concentration of incorporated Lys, measured while cooling and heating using differential scanning calorimetry (DSC). (b) DSC curves of MAPbBr_3_ around the phase transformation temperature, with and without incorporated Lys. The arrows indicate the direction of scanning.

The presence of Lys during growth has a noticeable effect on the crystal morphology **(Figure** 5a,b). As the amount of Lys in the solution increases, the crystals become more cubic-like, with clearer facets and sharper edges and corners. To examine these changes quantitatively, we looked at the diffraction peaks from different reflections of the MAPbBr_3_ samples **(Figure 5c**,**d**,**e)**: the relative intensities of the diffraction peaks from {110}, {120} and {222} decrease as the Lys incorporation levels increase. Since the intensity of the diffraction peaks is normalized to that of the {100} peak (the most intense diffraction peak in the MAPbBr_3_ diffractogram, see **Figure S1**), this decrease might also describe the increase in relative intensity of the corresponding {100} diffraction peak. Generally, the presence of Lys leads to an increase in intensity of the {100**}** reflection. Since the {100} family corresponds to the main facets of the cubic unit cell, this finding corroborates with the observed morphological changes depicted in **Figure 5a,b**.

**Figure 5.**
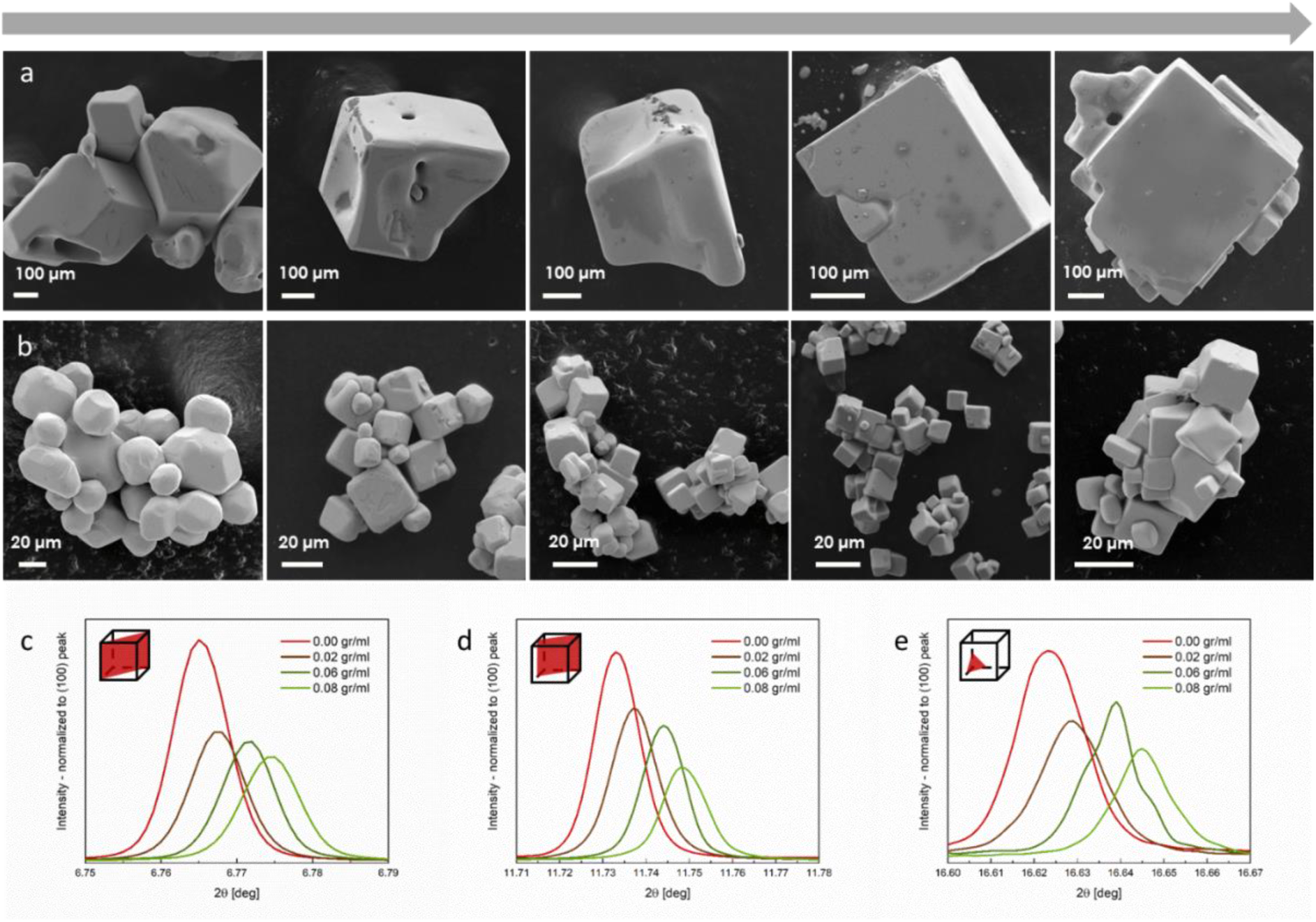
Morphology changes of MAPbBr_3_ crystals. HR-SEM micrographs of MAPbBr_3_ crystals grown in the presence of Lys in solution. (a) Slow growth; (b) Fast growth; The arrow indicates an increase in the concentration of incorporated Lys. (c)−(e) HR-PXRD reflections of different planes of slow-grown MAPbBr_3_: (c) {110}; (d) {120}; (e) {222}, grown with different amounts of Lys (λ=0.49599 Å). The intensity of the diffraction peaks are normalized to that of the corresponding {100} diffraction peak.

The fact that the incorporation of Lys induces higher lattice distortions than those of any of the other amino acids, combined with the effect of lattice parameter reduction, suggests the following incorporation mechanism: the two NH^3+^ groups of Lys probably replace two MA^+^ anions in the MAPbBr_3_ crystal, similarly to the replacement of carbonate groups in calcite by the COO^−^ group of Asp,^[69]^ and of Cu^2+^ in Cu^2^O.^[71]^ In the case of MAPbBr_3_ the Lys backbone acts as a “molecular bridge” that tightens the atoms together, thereby decreasing the lattice parameter and leading to the reduction in microstrain fluctuations **(Figure 2b,c)**. This bridging effect induces changes in MAPbBr_3_ thermal expansion coefficient as well as in its C-T transformation temperature.

In order to understand the crystallographic direction in which Lys molecules are incorporated into the MAPbBr_3_ crystal, a model was developed. This model takes into account the difference between the NH^3+^-NH^3+^ distance in Lys, and the MA^+^-MA^+^ distance in the crystal host, along with the ratio of stiffness coefficients for Lys and MAPbBr_3_ (for more details, see supporting information). We assume, that during fast growth, incorporation occurs along ⟨130⟩ direction, when two NH^3+^ groups substitute two adjacent MA^+^ ions, and O^-^ substitutes Br^-^ (**Figure 6b**). However, during slow growth, Lys becomes incorporated along ⟨110⟩ direction, and can either induce a Br^-^ vacancy (**Figure 6c**) or bend around it (**Figure 6d**). Note, that in both cases the incorporation direction is along the unit cell face, which can explain the passivation of the {100} planes upon Lys incorporation (**Figure 5**). These suggested mechanisms of Lys incorporation can be also responsible for the difference in incorporation levels between the two growth regimes (**Figure 1**c): since the incorporation occurs by attaching of Lys molecules to the {100} surface, the incorporation level should be higher for smaller crystals, which exhibit bigger surface area. According to **Figure 5** this effect is indeed confirmed experimentally.

**Figure 6.**
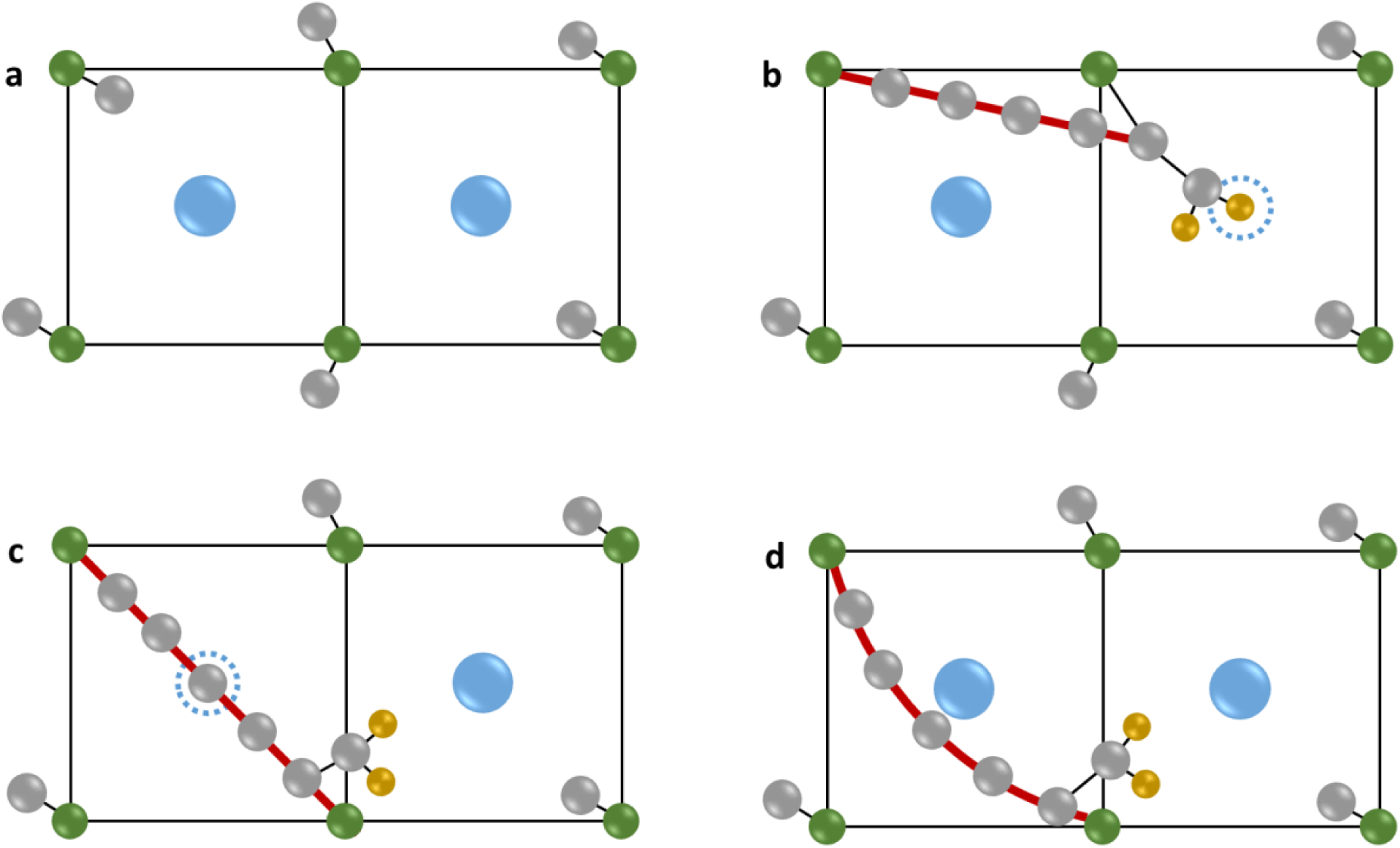
Different options for Lys incorporation. Schematic representation of two neighboring MAPbBr_3_ {100} planes. (a) Pure MAPbBr_3_, (b) MAPbBr_3_ with Lys incorporated along ⟨130⟩ during fast growth. (c+d) MAPbBr_3_ with Lys incorporated along ⟨110⟩ during slow growth, where Lys backcone (c) substitutes Br^-^, and (d) bends around Br^-^. The “molecular bridging” of the Lys backbone is marked in red. Color legend: N, green; C, gray; Br, blue; O, gold. The blue dotted line represents a Br^-^ vacancy. H atoms are not presented.

The remaining question to be answered is why incorporation along ⟨130⟩ is favoured during the fast growth, while incorporation along ⟨110⟩ is favoured during the slow growth? One possible explanation is the difference in the solubility of the precoursers over temperature. It is known that while Pb(OAc)_2_ solubility increses with temperature,^[84]^ the solubility of HBr decreases.^[85]^ In light of this, crystallization at low temperature and high Br^-^ solubility conditions promotes Lys to induce Br^-^ vacancies (**Figure 6**b) and allows for the substituon of the two adjacent MA^+^ ions for the NH^3+^ groups along ⟨100⟩ (while Lys backbone inclines along ⟨130⟩). However, at a higher temperature, Br^-^ solubility is lower and the induction of Br^-^ vacancies is suppresed. In the latter case, a Lys molecule substitutes MA^+^ ions located further apart, i.e. along ⟨110⟩.

Next, we investigated the effect of Lys incorporation on the optical properties of the host crystals. To this end we measured the diffusive reflectance (R) spectra of the powdered samples in the visible range (400–700 nm), using a spectrophotometer equipped with an integration sphere. Using the Kubelka-Munk theory, *i*.*e*.,^[86,87]^

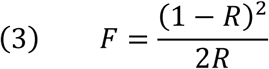

where *F* is proportional to the absorption coefficient, and based also on the fact that MAPbBr_3_ has a direct band gap,^[77,78]^ we plotted (*Fhv*)^2^ vs. the photon energy *hv*, taking the band gap as the x-intercept of the linear part of the plot^[88,89]^ (see **Figure S2**).

The incorporation of Lys into MAPbBr_3_ indeed induced an increase of more than 1% in its optical band gap (blueshift) **(Figure 7**). It is important to note that for a fixed amount of Lys, the change in the band gap is higher in the case of fast-grown samples, although the lattice distortions **(Figure 2b)** are lower. For the maximum amount of incorporated Lys in the slow-grown samples (0.2% at) the relative change in the band gap is only about 0.1%, that is six times lower than that of the fast-grown samples with similar amount of incorporated Lys. These findings may suggest that the factor governing the change in band gap is indeed the concentration of incorporated Lys (and the mechanism of incorporation), rather than the induced lattice distortions. This conclusion is consistent with the previously established understanding of the phenomenon^[72]^ whereby the incorporation of insulating organic molecules induces quantum confinement inside the bulk semiconductor crystals.

**Figure 7.**
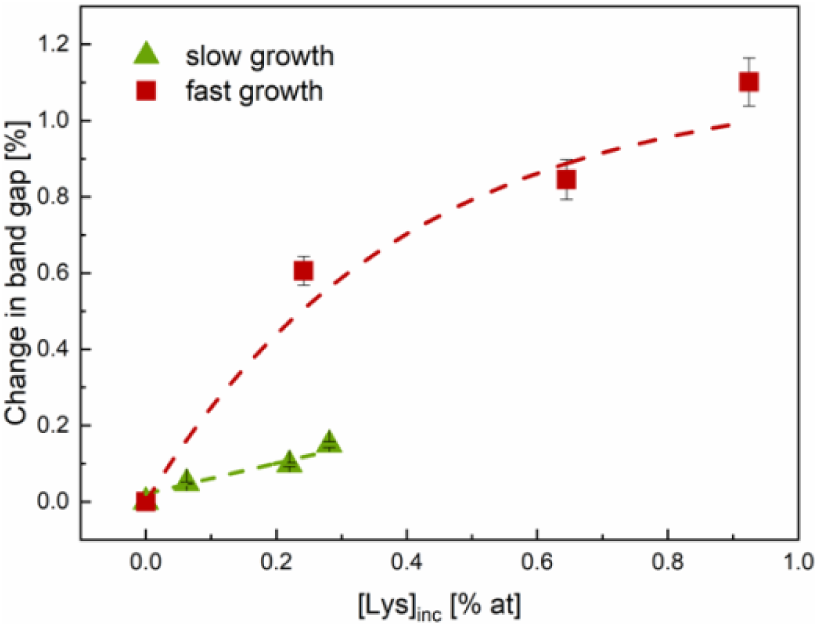
Band gap Changes. Change in the optical band gap of MAPbBr_3_ vs. concentration of incorporated Lys in the case of slow and fast growth.

The band gap of the MAPbBr_3_ crystals is formed due to an overlap between Pb-6s and Br-5p antibonding atomic orbitals, forming the valence band maximum.^[78]^ Removing Br^-^, as suggested by incorporation of Lys molecules along ⟨130⟩, should decrease this overlapping, hence increase the band gap. The more intensive increase of the band gap in the fast grown crystals manifests more substantial removal of Br^-^ anions in in the fast crystallization route. Incorporation of Lys molecules along ⟨110⟩ may occur without removal of Br^-^, when Lys molecule is bent. Apparently, such incorporation only slightly affects the electronic structure.

It is interesting to note that changes in the optical band gap can also originate from changes in hydrostatic volume.^[21,90]^ In our study, however, this is probably not applicable for two reasons. First, the band gap deformation potential (*i*.*e*., the change in band gap vs. the logarithm of the unit cell volume) is usually positive in HOIPs,^[90,91]^ as opposed to our results (see **Figure S3**). Secondly, a difference between the deformation potentials of the two synthetic routes for the same crystal was clearly seen in our study. Hence, we can conclude that in our case, similarly to that reported in the case of ZnO,^[72]^ changes observed in the band gap originate from quantum confinement effects, together with the subtitution of Br^-^ anions.

We also examined the effect of Lys incorporation on MAPbBr_3_ stability in water. Spesifically, we measured the kinetics of crystals solubility using an electrochemical approach, namely measurments of the solution impedance over time. The dissolution process of MAPbBr_3_ crystals results in the realease of free ions into the solution:

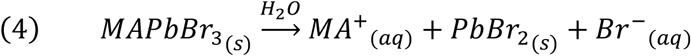

Hence, the impedance (*i*.*e*. resistivity) of the solution should decrese after the addition of MAPbBr_3_ crystals. We performed time-dependednt impedance measurments for several samples with different incoporation levels of Lys (**Figure S5**). The obtained curves were fitted according to equation (5):

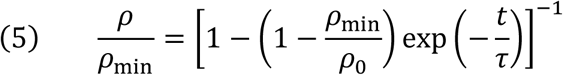

Where *τ* is the dacay time (the time needed to reach steady state), 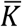 is the dissolution rate, 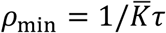 is the minimum resistivity (the plateau of the plot), and *ρ*_0_ is the maximum resistivity (prior to the addition of MAPbBr_3_). For more details, see the supporting information. **Figure 8**a presents the change in decay time of MAPbBr_3_ crystals upon the addition of Lys. In both slow- and the fast-grown samples, the decay time increases. However, it seems that the effect is much more significant in the slow regime. As was discussed above, the Lys molecules are situated on the {100} planes of the crystal. Hence, they hinder the ability of the ions to detach and/or attach to the crystal, and increase the time needed to achive steady state conditions. The dissolution rate, however, doubles upon Lys addition in the case of the fast grown MAPbBr_3_ crystals, and decrease by almost two fold in the case of the slow grown samples (**Figure 8**b). In the case of fast grown crystals, Lys molecules are incorporated in the crystal solely by subtitutions in the lattice. These subtitutions are weekly bound in the crystal, hence the incorporated molecules dissolve in water much easier. Conversely, during slow growth, CH_2_ groups of Lys interact with Br^-^ anions (as illustrated in **Figure 6**d). This interaction probably hinders the dissolution process, and decreases its rate. Overall, it is aparrent that the incorporation of Lys has a noticable effect on the dissolution kinetics of MAPbBr_3_ crystals. In order to enhance the stability of the hybrid perovskite crystals, amino acid incorporation during slow growth is required.

**Figure 8.**
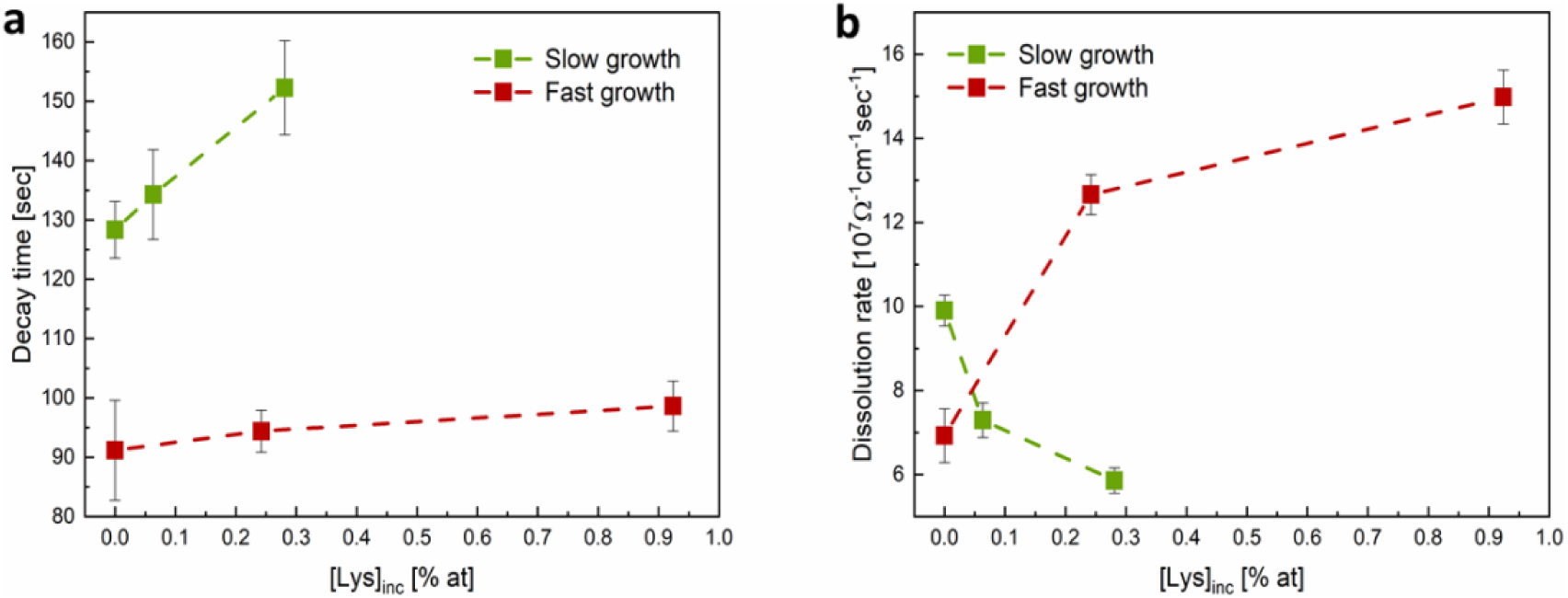
Effect on stabilty in water. (a) Decay time, and (B) dissolution rate of MAPbBr_3_ in water, as was measured using time-dependednt impedance measurments.

## Conclusion

The present study shows, for the first time, that the possibility of utilizing amino acid incorporation in order to tune the physical properties of crystalline hosts is not restricted to purely inorganic materials. Here we demonstrated a similar phenomenon with a hybrid semiconductor whose lattice includes both organic and inorganic components.

Incorporation of Lys molecules into the MAPbBr_3_ lattice depends on the route of crystal growth. At high temperatures (slow crystal growth) Lys incorporation occurs along ⟨110⟩ direction and results in a significant lattice contraction and slight band gap change. However, at lower temperatures (fast crystal growth) Lys becomes incorporated along ⟨130⟩ and causes opposite effects in the crystals, namely strong lattice contraction accompanied with considerable band gap increase. This observation can be explained by the tendency of Lys molecules to induce lattice shrinkage in MAPbBr_3_ upon incorporation and decrease in HBr solubility with increasing temperature. We support this interpretation with experimental finding of a decrease in the thermal expansion coefficient of MAPbBr_3_ upon Lys incorporation. Moreover, Lys incorporation has a distinct effect on the stability of MAPbBr_3_ in water. While the time required to achieve steady state increases upon the incorporation of Lys in both growth regimes, Lys incorporation during slow growth decreases MAPbBr_3_ dissolution rate by ∼40%.

Recently, the addition of amino acids during MAPbBr_3_ synthesis was shown to passivate the crystal surface, facilitating the formation of nanocrystals thereby improving devices performence.[92,93] In our study, we showed that not only do the amino acids passivate the MAPbBr_3_ crystal surface, but they also become incorporated into its lattice, thus, alter the optical and thermal properties of the perovskite. Another cardinal outcome of the incoporation is the finding that it enhances the stability of the host MAPbBr_3_ crystal under humid conditions.

We believe that these new discoveries can shed light on the interaction between amino acids and different crystalline systems. The mechanism of Lys incorporation into a non-oxide hybrid crystal, as shown here, differs from that reported previously, and can provide a new method for fine tuning of band gap and improved stability of hybrid systems. This latter point will facilitate the commercializing of halide perovskites. Moreover, the observed structural changes induced by such incorporation provide additional evidence for the intriguing interaction between the organic amino acids and the inorganic (or hybrid) crystals, which is still far from being completely understood.

## Methods

### MAPbBr_3_ Crystal Growth

Crystals were synthesized according to the method described by Potlatch & Weber.^[17]^ In a typical process, we prepared a Pb^+2^ solution by dissolving lead acetate trihydrate (2 g Pb(OAc)_2_*3H_2_O; Merck) in concentrated hydrobromic acid (10 ml 48% HBr; Sigma Aldrich). Different amounts of all the 20 common L-amino acids were added to the Pb^+2^ solution: L-Alanine (Ala), L-Arginine (Arg), L-Asparagine (Asn), L-Aspartic acid (Asp), L-Cysteine (Cys), L-Glutamic Acid (Glu), L-Glutamine (Gln), Glycine (Gly), L-Histidine (His), L-Isoleucine (Ile), L-Leucine (Leu), L-Lysine (Lys), L-Methionine (Met), L-Phenylalanine (Phe), L-Proline (Pro), L-Serine (Ser), L-Threonine (Thr), L-Tryptophan (Trp), L-Tyrosine (Tyr) and L-Valine (Val). Each solution was stirred until dissolution was complete. Meanwhile, we prepared a 1:1 (v/v) stock solution of MA^+^ in HBr by adding a methylammonium hydroxide (MAOH) solution (40% in H_2_O; Merck) dropwise into concentrated HBr, while stirring in an ice/water bath. MAPbBr_3_ precipitation was achieved simply by the addition of equimolar volumes of MA+ solution into the Pb^+2^ solution. This was done via two routes: fast synthesis, in which we added the MA^+^ to the Pb^+2^ solution at room temperature while stirring, and slow synthesis, in which we added the MA^+^ to a hot Pb^+2^ solution (in an oil bath at 95°C) and allowed the mixture to cool down naturally. In both cases, after precipitation the orange-hued crystals were filtered, washed with acetone, and dried in air.

### Chemical Analysis

Amino acid analysis (AAA) was carried out at AminoLab, Rehovot, Israel. Samples were sonicated in concentrated HBr and then washed several times with HBr and acetone to remove any remaining Lys from the surface. To ensure that no Lys was left after this process, we mixed a pure sample (without Lys) in a Lys-containing HBr solution and subjected it to the same treatment. Samples were filtered and dried in air. Each sample was then dissolved in 0.1% HCl and injected into a column to separate the Lys from the MA^+^ cations that had originated from the crystals. This was followed by post-column derivatization with ninhydrin and detection in a standard UV-Vis detector.^[94,95]^

### Structural Characterization

Samples were subjected to high-resolution powder X-ray diffraction (HR-PXRD) in ID22 at the European Synchrotron Radiation Facility (ESRF) in Grenoble, France, at a monochromatic radiation of 0.49599 Å (corresponding to 25 keV). Each sample was transferred to a 0.9-mm glass capillary and scanned three times at a fast rate (10 deg/min) at three different positions, while being rotated. This setup makes it possible to avoid beam damage and texture influences. We used the Rietveld refinement method in GSAS-II software for data analysis and lattice parameter calculations^[79]^. Gnuplot software was used for peak fitting.

### Morphology

Crystal morphology was observed by using a Zeiss Ultra Plus high-resolution scanning electron microscope (HR-SEM).

### Thermal Properties

We measured the lattice parameter of each sample at different temperatures (290°K, 275°K, 250°K, 230°K) in ID22 at ESRF, as described above. Cooling was achieved with a cold nitrogen gas blower. The lattice expansion coefficient was calculated according to Equation (2):

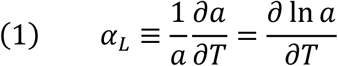

i.e., the slope of ln(a) (when a is the measured lattice parameter) vs. the absolute temperature. To evaluate the phase transformation temperature, we used DSC. Each sample was scanned from room temperature to −80°C and back, at a rate of 5°C/min (DSC 3+; Mettler Toledo). The transformation temperature was taken as the temperature at onset of the peak.

### Optical Properties

We measured the diffusive reflectance (*R*) of the powders in the visible range (400–700 nm) using a spectrophotometer equipped with an integration sphere (Cary 5000, Agilent). According to the Kubelka-Munk theory^[86,87]^:

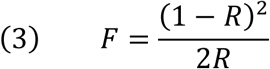

*F* is proportional to the absorption coefficient. Hence, based on the fact that MAPbBr_3_ has a direct band gap,^[77,78]^ *(Fhv)*^*2*^ was plotted vs. the photon energy *hv*, and the band gap was taken as the x-intercept of the linear part of the plot.^[88,89]^

### Stability Measurements

We measured the kinetics of MAPbBr_3_ crystals solubility in water using a simple electrochemical set-up. A standard reference electrode was immersed in DI water while stirring. The impedance of the solution was measured over time, at a constant frequency of 100 kHz. At a known time, a known amount of crushed MAPbBr_3_ crystals was added to the solution, and a significant decrease in the resistivity of the solution was observed. The measurement was carried out until reaching a plateau of the impedance vs. time curve.

## Acknowledgements

We would like to acknowledge ID22 beamline at the ESRF (Grenoble, France) and Dr. C. Dejoie and Dr. W. Mashikoane for helping in collecting the powder diffraction data. Additionally, we would like to thank G. Kozyukin from the Israel Institute of Metals (IIM) for his help with the electrochemical measurements. This research was supported by a Grant from the GIF, the German-Israeli Foundation for Scientific Research and Development No. I-1512-401.10.

## Data availability

The data that support the findings of the study are available from the corresponding author upon reasonable request.

## Author contributions

A.L. have performed the experiments in this work. A.K. has developed the model. A.L., I.P., E.S., J.F., A.K., and B.P. have participated to the redaction of the manuscript. All authors have read and approved the manuscript. B.P. provided supervision of the research.

## Competing interests

The authors declare no competing financial interests.

## Notes

### Competing Interest Statement

The authors have declared no competing interest.

### Summary of Updates

references corrected

